# Low-fidelity simulator selection for multi-fidelity inference of epithelial vertex models

**DOI:** 10.64898/2026.10.29.762786

**Authors:** Mohammad Amini, Jack L. Jennings, Wei W. Xing, Alexander G. Fletcher

## Abstract

Vertex models link cell division, neighbour exchange, and intercellular forces to epithelial tissue form through parameters such as junctional line tension and perimeter contractility. Bayesian inference of these parameters from data quantifies their uncertainty, but for stochastic models of proliferating tissues it requires many costly simulations. Multi-fidelity simulation-based inference reduces this cost by training on many runs of a cheaper, less accurate (‘low-fidelity’) simulator before refining with a few full (‘high-fidelity’) runs. Low-fidelity versions can be constructed in several ways, for example by stopping simulations early, coarsening the timestep, or averaging fewer replicates, but it is unclear which to choose before committing to costly training. To address this, we introduce PRISM (Paired Ranking Index for Simulator Matching), a screen that compares each candidate with the full simulator on paired runs, scoring agreement in cell-packing statistics against cost. For a vertex model of a proliferating tissue, PRISM flags stopping early as unreliable before training; once trained, this choice can produce confidently wrong parameter estimates. By contrast, a coarser timestep, within the range tested, is cheap and accurate; while fewer replicates are accurate but too costly. Using a screened low-fidelity simulator, multi-fidelity inference outperforms approximate Bayesian computation and neural-network inference trained on full simulations alone, at matched cost and with well-calibrated uncertainty. The same pattern recurs for a second vertex model fitted to published *Drosophila* pupal-wing imaging data, although there stopping halfway proves safe. Together, our findings suggest that which approximations are safe depends on the model, and that screening them before training makes multi-fidelity inference of tissue mechanics practical.

**Author summary:** As tissues grow and take shape, their cells divide, push and pull on their neighbours, and swap places. Computer models can simulate this cell by cell, and fitting them to measurements can reveal mechanical properties, such as the tension along cell-cell junctions, that cannot be measured directly. However, fitting is slow, because the models include randomness and so each candidate must be simulated many times. A promising shortcut is to do most of the work with cheap, rough versions of the simulation and only a little with the full version. The difficulty is knowing in advance which rough versions can be trusted. We develop a quick check, which we call PRISM, that compares cheap versions with the full simulation before any lengthy fitting begins. Not all shortcuts are equal: stopping simulations early can give confidently wrong answers, taking larger timesteps stays reliable, and averaging fewer runs is accurate but saves too little time. Guided by PRISM, the shortcut approach gives better estimates than standard methods for the same computing effort. The same checks worked for a second model fitted to real images of a fruit fly wing, although there a different shortcut proved safe, underlining the value of checking first.

## Introduction

Epithelial tissues are remodelled throughout animal development as cells grow, divide, and exchange neighbours, generating collective flows and changes in tissue shape [1]. Vertex models represent the apical surface of such a tissue as a dynamic tessellation of polygons, linking cell-scale mechanics to tissue-scale morphogenesis [2, 3]. Their parameters, such as the tension along cell-cell junctions, represent physical properties that are difficult to measure directly, and changes in such properties can have clear biological consequences. For example, Park and colleagues showed that a cell shape index predicted by vertex models distinguishes the fluid-like, unjammed state of asthmatic airway epithelium from the solid-like state of healthy tissue [4]. Estimating these parameters from tissue data, together with their uncertainty, is therefore an important goal. However, it is computationally expensive, because the cell divisions and neighbour exchanges that give vertex models their explanatory power must be simulated to completion many times before simulated tissues can be compared with experimental data [5].

Existing approaches to estimating the parameters of vertex models fall into two broad classes, depending on whether or not the model is simulated forward in time. The first class infers mechanical forces directly from static tissue geometry. For example, Chiou and colleagues recovered junctional tensions and cell pressures from force balance on segmented cell arrays [6], and Ishihara and Sugimura extended this approach to Bayesian estimation from *Drosophila* images [7]. More recent work fits vertex-model energies to segmented images by assuming that the tissue is close to mechanical equilibrium [8, 9], or through gradient-based optimisation of a differentiable vertex model [10]. These methods are well suited to static or smoothly varying vertex models, but cannot readily be applied to models of proliferating tissues, in which cell division and neighbour exchange introduce randomness and abrupt changes in tissue topology, as implemented in platforms such as Chaste [11].

The second class, which we follow here, simulates the model forward in time and compares its output with data. We focus on the widely used energy-based vertex model of Farhadifar and colleagues, in which junctional line tension (Λ) and perimeter contractility (Γ) determine how cells pack together (Fig 1A) [12]. In previous work, we showed that numerical implementation choices, such as the integration timestep, affect the quantitative predictions of such models even when their mechanical parameters are held fixed [13]. We subsequently used approximate Bayesian computation (ABC) to estimate Λ and Γ for a vertex model of a proliferating tissue implemented in Chaste [14]. That study found that reliable estimates required summary statistics averaged over repeated simulations of each tissue, and the analysis required 10^5^ parameter samples, each simulated ten times. It defines the model, summary statistics, and inference task that we adopt here, and illustrates the computational cost that motivates the present work. Similar costs arise when calibrating other cell-based models, including cellular Potts [15], agent-based [16], spatial-stochastic [17], and subcellular-element [18] models. In each case a single simulator is used throughout, and the question of whether a cheaper approximation could safely replace it for part of the computation is not addressed.

**Fig 1.**
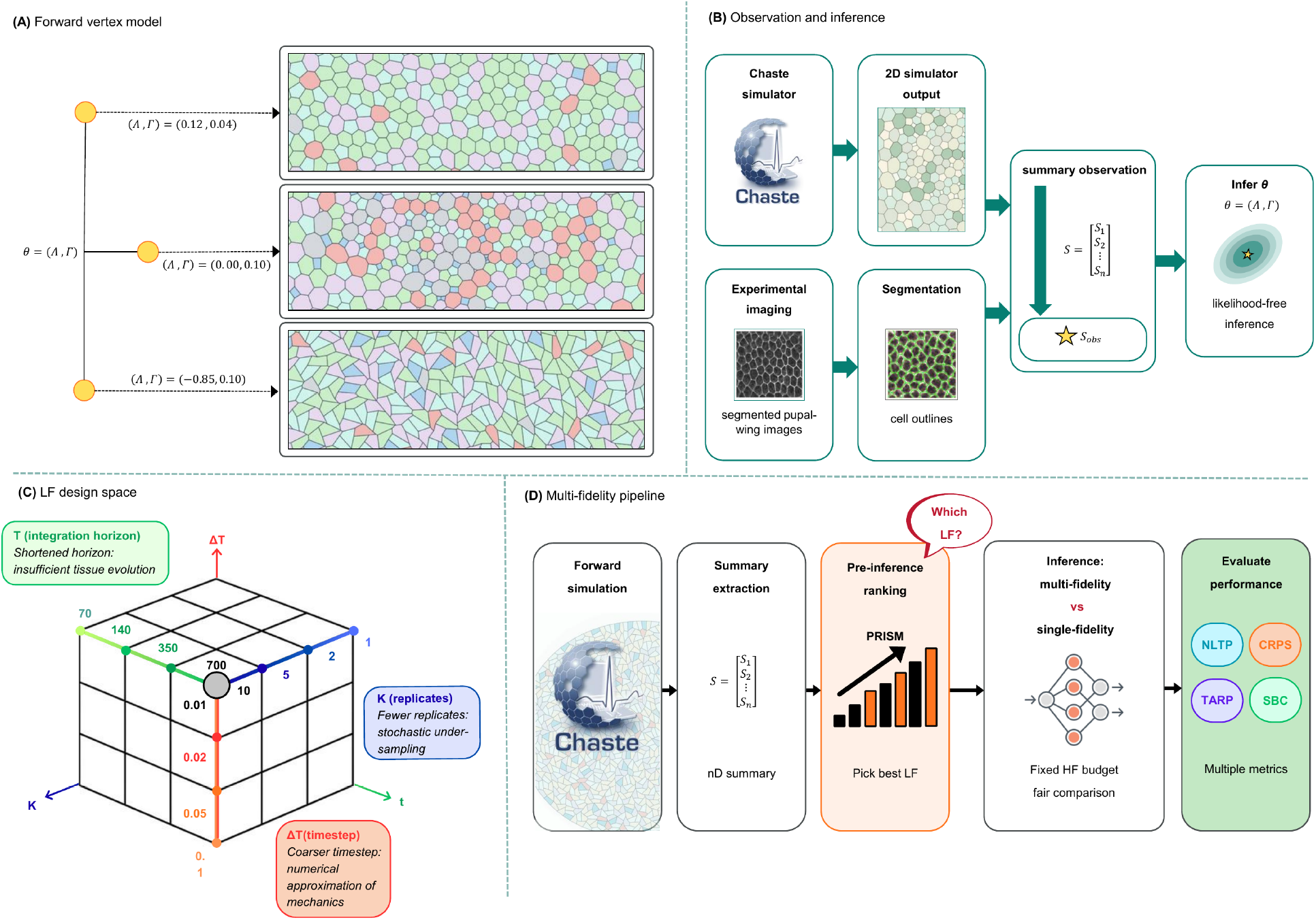
PRISM screens low-fidelity (LF) approximations of a vertex model of a proliferating epithelial tissue before multi-fidelity (MF) inference. (A) Simulated epithelial packing at three values of the mechanical parameters *θ* = (Λ, Γ). (B) Six summary statistics of cell packing, computed from simulated or segmented tissue, are used to infer *θ*. (C) Nine LF candidates, obtained by shortening the simulation duration *T* , coarsening the timestep Δ*T* , or reducing the number of averaged replicates *K* relative to the high-fidelity (HF) reference (*T*_end_ = 700, Δ*T* = 0.01, *K* = 10). (D) PRISM screening precedes MF and HF-only inference under a matched computational budget; the resulting posteriors are assessed for accuracy and calibration (see *Materials and Methods*).

Inference of this kind is often described as ‘likelihood-free’ or ‘simulation-based’. Since each simulation is stochastic, and its output must first be reduced to a small number of summary statistics (for example, the mean area of cells with each number of neighbours), the probability of observing particular data given a set of parameter values cannot be written down explicitly [19]. Instead, inference proceeds by comparing simulated and observed summary statistics. The classical approach, sequential Monte Carlo ABC (SMC-ABC), accepts parameter values whose simulated summary statistics fall within a tolerance of the observed data, and progressively tightens this tolerance [20–22]. More recent neural simulation-based inference (SBI) methods instead use pairs of parameter values and simulated outputs to train a neural network that approximates the posterior distribution of the parameters. Examples include neural posterior estimation (NPE) and its sequential variant (SNPE), which concentrates later simulations in the most plausible regions of parameter space [23–27]. Benchmark comparisons show that neural methods can outperform ABC, although their relative performance depends on the metric used [28], and the sbi toolkit has made them widely accessible [29, 30]. We use SMC-ABC, NPE, and SNPE as baselines in this study. Each relies solely on the full, or ‘high-fidelity’ (HF) simulator, so every simulation used for training or acceptance incurs its full cost.

Multi-fidelity methods reduce this cost by combining many runs of a cheaper, less accurate ‘low-fidelity’ (LF) simulator with a small number of HF runs, an idea that has been widely used in uncertainty quantification, inference, and optimisation [31]. In multifidelity Monte Carlo estimation, the benefit of an LF model depends on its correlation with the HF model and on its relative cost, and these two quantities can be used to choose which of several LF models to include [32]. Prescott and Baker brought these ideas to likelihood-free inference, developing multifidelity ABC algorithms in which cheap LF simulations are used to make early decisions about whether to accept or reject each proposed parameter value, with HF simulations run only where needed [33, 34]; this approach has since been combined with multilevel Monte Carlo [35].

Within neural SBI, the idea is more recent. Krouglova and colleagues introduced multi-fidelity neural posterior estimation, in which a neural network is first trained on LF simulations and then refined on HF simulations [36]; this approach has since been extended to multilevel training [37] and to transfer learning between related tasks [38]. We refer to such neural approaches collectively as multi-fidelity SBI (MF-SBI). However, both the ABC-based and neural approaches assume that a suitable LF simulator is already available: the former optimise how a given LF simulator is used, while the latter give no guidance on how accurate or how cheap it needs to be. For cell-based models, several LF simulators can typically be constructed from the same HF model, for example by stopping simulations early, using a coarser integration timestep (taking care with numerical stability), or averaging over fewer stochastic replicates. These choices are not equivalent: an approximation that is cheap but fails to reproduce the features of the data on which inference relies may produce misleading parameter estimates. Yet there is currently no general procedure for choosing among candidate LF simulators for simulation-based inference before committing to costly training.

To address this, we introduce PRISM (Paired Ranking Index for Simulator Matching), a screening protocol that compares each candidate LF simulator with the HF simulator before any neural network is trained. PRISM uses paired LF and HF simulations, run at the same parameter values and random seeds, to assess how well each candidate reproduces the HF summary statistics, and weighs this against its computational cost. Candidates can then be classified as cheap and accurate, accurate but too costly, or cheap but unreliable, and only the first group need be taken forward for training. Since PRISM requires only paired simulations and their summary statistics, it can in principle be applied to any stochastic simulator for which several reduced-cost versions are available.

We apply PRISM to the vertex model case study of [14], screening nine LF simulators constructed by shortening the simulation duration, coarsening the timestep, or reducing the number of averaged replicates (Fig 1). We find that these approximations are not interchangeable. Stopping simulations early is flagged as unreliable before training and, once trained, can produce posteriors that place less probability on the true parameters than the prior; coarsening the timestep, within the range tested, is cheap and accurate; and reducing the number of replicates preserves accuracy but is too costly to be useful under a limited budget. Using a screened LF simulator, MF-SBI gives more accurate posteriors than SMC-ABC, NPE, and SNPE at a matched computational budget equivalent to 1,000 HF simulations, while using only 500 HF simulations, and its calibration at the selected configuration is confirmed across 20 independent test observations. Finally, we apply the same approach to a second vertex model of a non-proliferating tissue, fitted to published imaging data from the *Drosophila* pupal wing [9]. The same three classes of candidate emerge, although here stopping simulations halfway proves both cheap and accurate. This indicates that which approximations are safe depends on the model, and underlines the value of screening candidates before training. Fig 1 summarises this workflow.

## Results

We first applied PRISM to nine candidate LF simulators for the vertex model of a proliferating tissue, before any neural network was trained. We then performed multi-fidelity inference with every candidate, to test whether the screening results predict the accuracy of the resulting posteriors, and checked the calibration of the posterior for the selected configuration. Finally, we repeated screening and inference for a second vertex model, fitted to published imaging data from the *Drosophila* pupal wing. Definitions of all screening metrics and posterior measures are given in *Materials and methods*.

### Screening identifies cheap and accurate low-fidelity simulators before training

We constructed nine LF simulators from the HF simulator, which averages summary statistics over *K* = 10 realisations with *T*_end_ = 700 and Δ*T* = 0.01 (Table 3). Three use a single realisation with a shorter simulation duration (*T*_70_, *T*_140_, *T*_350_), three use a single realisation with a larger timestep (Δ*T*_0.02_, Δ*T*_0.05_, Δ*T*_0.10_), and three use HF settings but average over fewer realisations (*K*_1_, *K*_2_, *K*_5_). We compared each candidate with the HF simulator at 20 parameter values spread across Region B, using ten common random seeds at each (*Materials and methods*).

Shortening the simulation duration had the largest effect on agreement with the HF simulator (Fig 2A–C). The two shortest durations, *T*_70_ and *T*_140_, gave negative values of *R*^2^ (*−*0.88 and *−*1.26; Fig 2B), meaning that their summary statistics were further from the HF outputs than the mean HF output was, and they also showed the largest differences in distribution (Fig 2C). *T*_350_ gave a positive *R*^2^, but the weakest correlation with the HF simulator in any summary statistic among the remaining candidates (*ρ*_min_ = 0.67; Fig 2A). By contrast, all three timestep candidates and all three replicate candidates agreed closely with the HF simulator, with *R*^2^ *≥*0.98 and *ρ*_min_ *≥*0.95.

**Fig 2.**
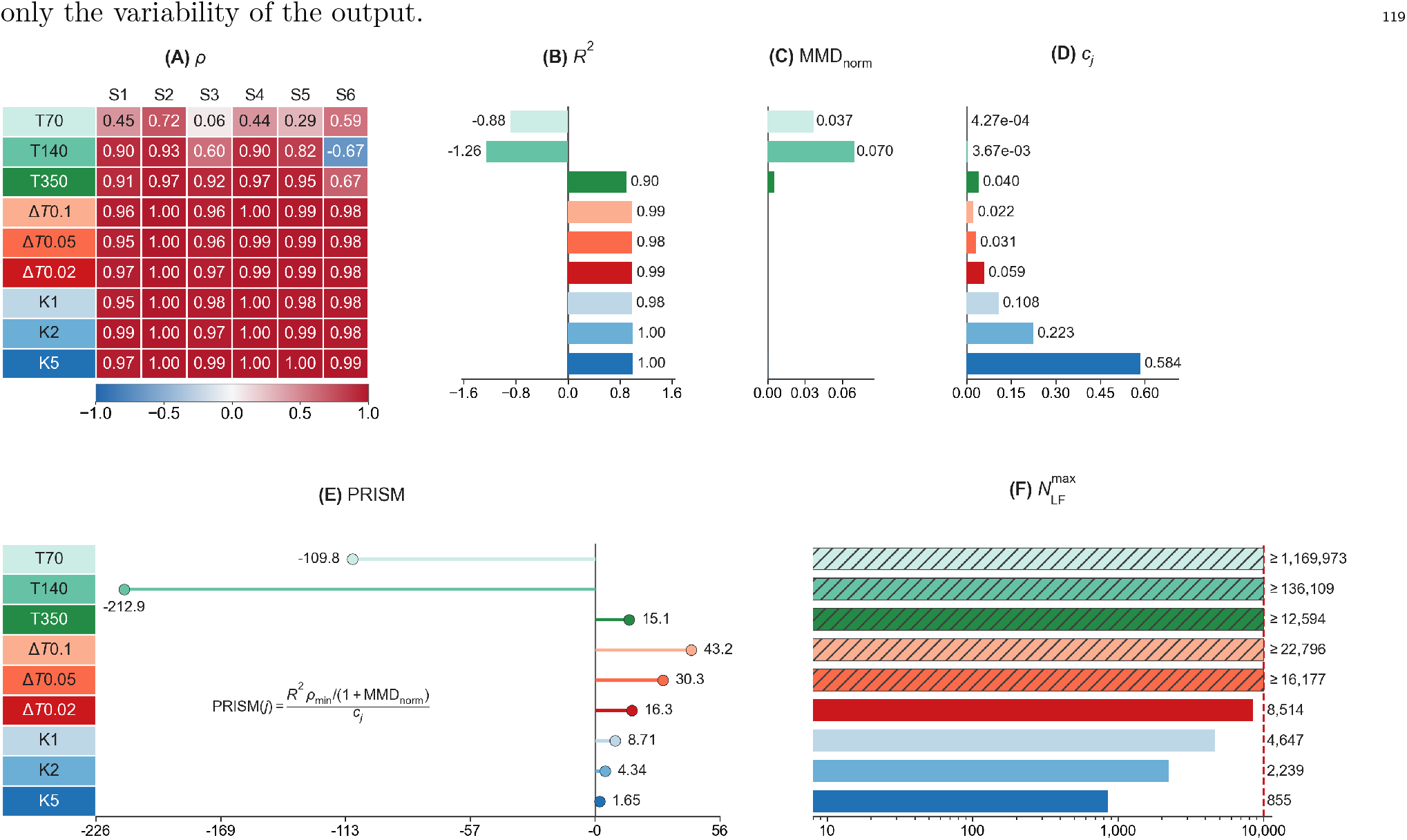
Screening of nine LF simulators for the vertex model of a proliferating epithelial tissue. Each candidate was compared with the HF simulator at 20 parameter values in Region B, using ten common seeds at each. (A) Pearson correlation *ρ* between LF and HF values of each summary statistic (S1–S6). (B) Pooled coefficient of determination, *R*^2^; only *T*_70_ and *T*_140_ are negative. (C) Normalised maximum mean discrepancy, (MMD_norm_), which is largest for the same two candidates. (D) Relative cost, *c*_*j*_. (E) PRISM score (Eq (8)). (F) Largest number of LF simulations affordable within a budget of 1,000 HF-equivalent units after reserving 500 HF simulations, 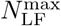 (Eq (9)); hatched bars exceed the axis range.

These differences can be understood from the timing of the simulations. With a mean cell-cycle duration of 20 time units, the seven rounds of division take on average around 140 time units (*Materials and methods*). Simulations stopped at *T*_end_ = 70 therefore end part-way through proliferation, while those stopped at *T*_end_ = 140 end at around the time that division is completing, before the tissue has relaxed. In both cases, the cell packing captured by the summary statistics has not yet formed. A larger timestep, in contrast, changes only the numerical resolution of a complete simulation, and averaging over fewer realisations changes only the variability of the output.

The candidates also differed greatly in cost (Fig 2D). Each replicate candidate simulates the full duration at the HF timestep; therefore, its cost is roughly proportional to *K*, ranging from *c*_*j*_ = 0.11 for *K*_1_ to 0.58 for *K*_5_. Within our budget of 1000 HF-equivalent units, with 500 reserved for HF simulations, this leaves room for at most 4647 LF simulations with *K*_1_ and 855 with *K*_5_ (Fig 2F). The timestep candidates were several times cheaper (*c*_*j*_ = 0.022–0.059), and every timestep and duration candidate except Δ*T*_0.02_ (8,514) could afford the largest LF budget we considered (10000).

Combining agreement and cost, the PRISM score was highest for Δ*T*_0.10_ (43.2) and Δ*T*_0.05_ (30.3), and negative for *T*_70_ (*−*110) and *T*_140_ (*−*213), which failed the sign test (Fig 2E). Plotting the accuracy term against cost separates the candidates into three groups (Fig 3A). The timestep candidates are cheap and accurate (efficiency-optimal); the replicate candidates are accurate but costly (affordability-bound); and the duration candidates are inaccurate regardless of cost (fidelity-bound). In practice, only the efficiency-optimal group would be taken forward to inference. Here, to test whether the screen predicts the outcome of inference, we trained with all nine candidates.

**Fig 3.**
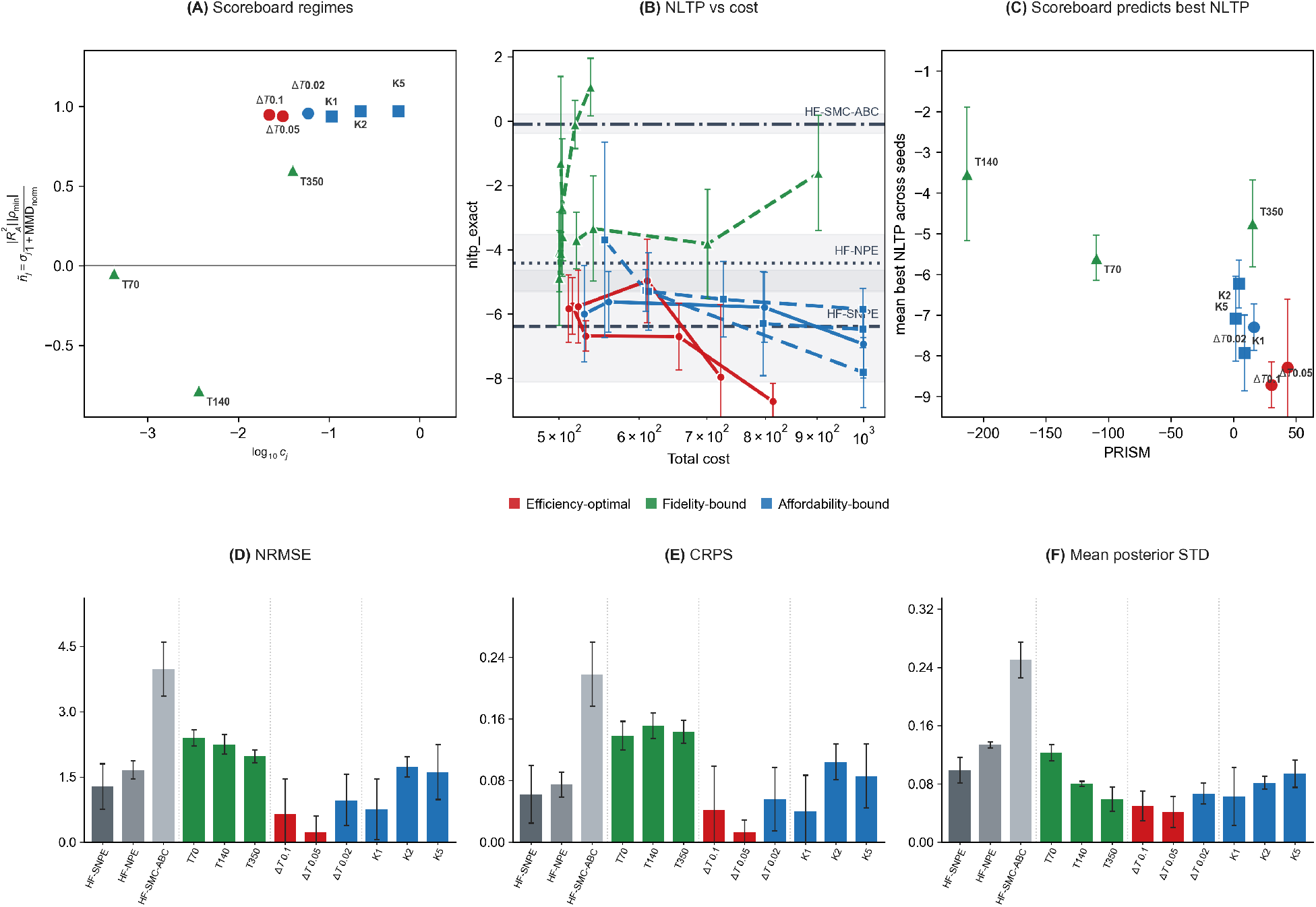
Screening results predict posterior accuracy after multi-fidelity training. (A) Accuracy term of the PRISM score plotted against relative cost, separating efficiency-optimal (red), fidelity-bound (green), and affordability-bound (blue) candidates. (B) Negative log target probability (NLTP) at *θ*^*^ against total cost in HF-equivalent units, for MF-SBI with each candidate across LF budgets; horizontal lines show the HF-only baselines, each using 1000 HF simulations. Lower NLTP indicates more posterior probability at *θ*^*^. (C) Best NLTP after training, over LF budgets, against PRISM score. (D–F) Normalised root mean square error (NRMSE), continuous ranked probability score (CRPS), and mean posterior standard deviation (SD) for each method, at the largest affordable LF budget. All values are mean *±* SD over five repeats; the error bars show that standard deviation. MF-SBI used 500 HF simulations throughout.

Simulation failures were common for all candidates except the shortest durations, with failure rates of up to 55% per seed (Fig 8B), and HF failure rates varied widely across the screening parameter values (Fig 8A). A run fails when the mesh forms a self-intersectionand the solver aborts, or when it stops making progress and is stopped. In this screen the recorded failures were self-intersections and stalled runs. A *K*_2_ output is the mean of two realisations and a *K*_5_ output the mean of five, and that output is counted as failed if any one realisation fails. These candidates therefore fail more often than *K*_1_: 44% and 52% of screening runs, against 28% for *K*_1_.

### Screening results predict posterior accuracy after training

We next performed multi-fidelity inference with each candidate, using 500 HF simulations and LF budgets of up to 10000 simulations (capped at 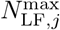), and compared the results with three HF-only baselines: SMC-ABC, NPE, and SNPE, each using 1000 HF simulations. Every configuration was repeated five times (*Materials and Methods*).

The groups identified by screening matched the outcome of inference (Fig 3B,C). The fidelity-bound candidates gave the least accurate posteriors of any method. For *T*_140_, the NLTP at the largest LF budget exceeded that of the prior, meaning that the trained posterior placed less probability on the true parameters than the prior did. The uniform prior has NLTP *≈ −*1.13. All five *T*_140_ repeats lay above that value, with mean NLTP about 1.1. *T*_350_, which passed the sign test but had the weakest single-statistic correlation, also failed to give a competitive posterior, confirming that a positive correlation with the HF simulator is not sufficient on its own. The affordability-bound replicate candidates became competitive with the weaker HF-only baselines once enough of the budget had been spent on them, but did not match the efficiency-optimal candidates.

The efficiency-optimal candidates gave more accurate posteriors than every other candidate and every HF-only baseline. With Δ*T*_0.05_, MF-SBI reached NLTP *≈−*8.6 at a total cost of roughly 800 HF-equivalent units, compared with *−*6.6 for SNPE, the best HF-only baseline, *−*5.4 for NPE. SMC-ABC has no parametric density. A Gaussian kernel density on its 300 particles, using Scott’s bandwidth, gives NLTP *≈ −*1.1, close to the prior and much higher than the NPE and SNPE values. Δ*T*_0.10_ reached similar accuracy at a similar cost. Within this group, the order of the PRISM scores was not preserved: Δ*T*_0.10_ had the higher score but Δ*T*_0.05_ gave the more accurate posterior (Fig 3C). This is consistent with how we intend the score to be used, to identify the group of candidates worth training rather than to rank them precisely.

The other measures of posterior accuracy gave the same picture (Fig 3D–F). MF-SBI with Δ*T*_0.05_ gave the lowest CRPS and NRMSE of any method, followed by Δ*T*_0.10_, and both gave narrower posteriors than the HF-only baselines. SMC-ABC gave the broadest posteriors of the three baselines. Within the efficiency-optimal group, therefore, the screen identifies which candidates are worth training, and posterior measures then identify the best configuration.

Posterior samples for the primary observation show these differences directly (Fig 4). SMC-ABC gave a broad posterior centred away from *θ*^*^. NPE and SNPE recovered the negative correlation between Λ and Γ that we observed previously [14], but retained considerable probability away from *θ*^*^. MF-SBI with Δ*T*_0.05_ gave a narrower posterior concentrated around *θ*^*^. In contrast, MF-SBI with *T*_140_ gave a narrow posterior that largely excluded *θ*^*^ (S1 Fig), illustrating how an unsuitable LF simulator can lead to confident but wrong estimates.

**Fig 4.**
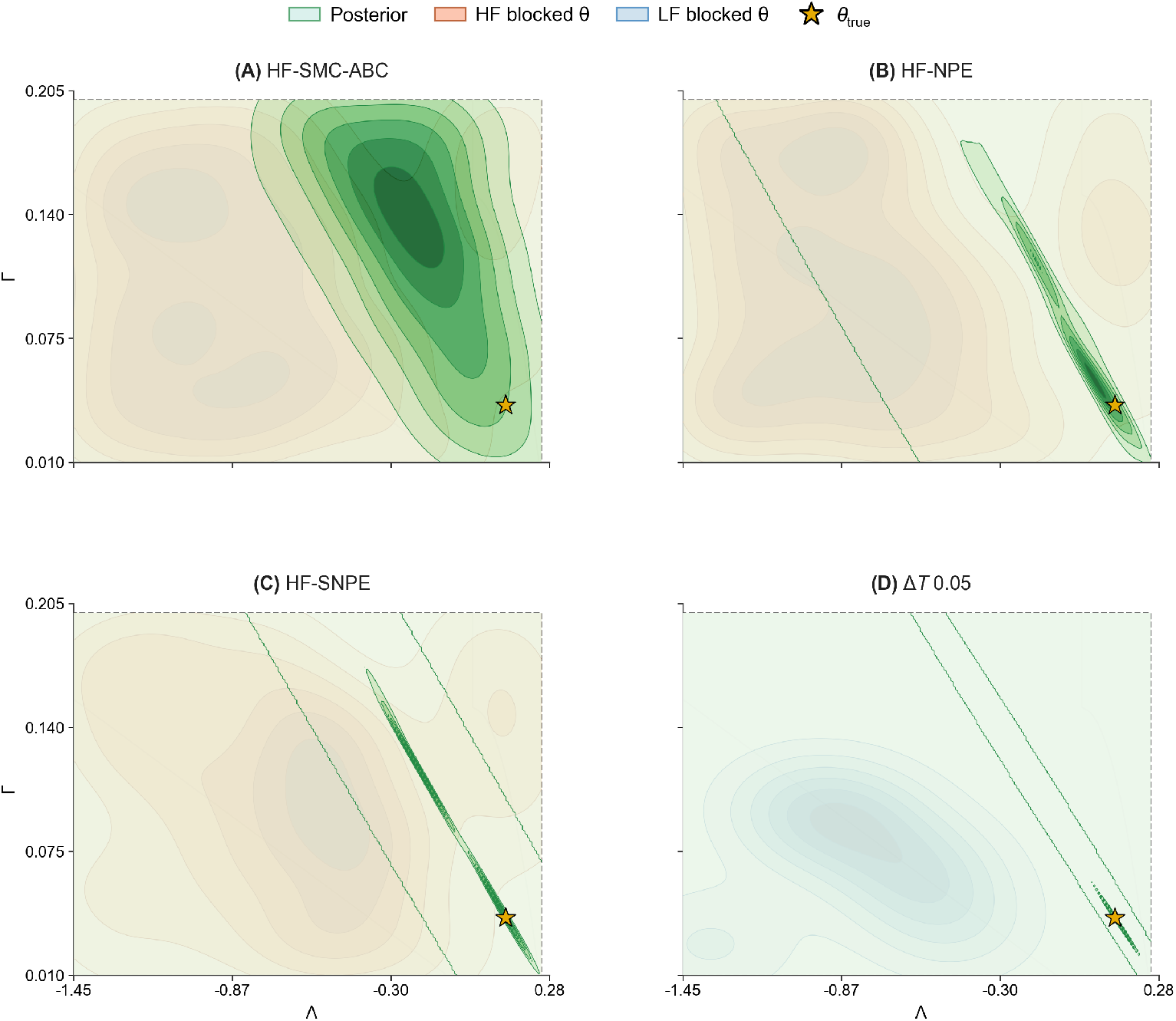
Posteriors for the primary observation, *θ*^*^ = (0.12, 0.04), **at a matched budget**. (A) SMC-ABC, (B) NPE, and (C) SNPE, each using 1000 HF simulations. (D) MF-SBI with the Δ*T*_0.05_ LF simulator, using 500 HF and 10000 LF simulations. Stars mark *θ*^*^. The shading is a density of the points of the same colour. Green shows posterior samples. Orange shows parameters at which a high-fidelity simulation failed (HF blocked *θ*). Blue shows parameters at which a low-fidelity simulation failed (LF blocked *θ*).

### The selected configuration gives well-calibrated posteriors

Accuracy for a single observation does not establish that the uncertainty reported by the posterior is reliable. We therefore assessed the calibration of the selected configuration, MF-SBI with Δ*T*_0.05_, 10000 LF and 500 HF simulations, using simulation-based calibration (SBC) [39] and Tests of Accuracy with Random Points (TARP) [40] on 20 further observations chosen from 200 parameters drawn uniformly from the prior box at which all ten high-fidelity replicates finished (Fig 5). We carried out this check only for the selected configuration, since repeating it for every candidate and budget would have dominated the computational cost of the study.

**Fig 5.**
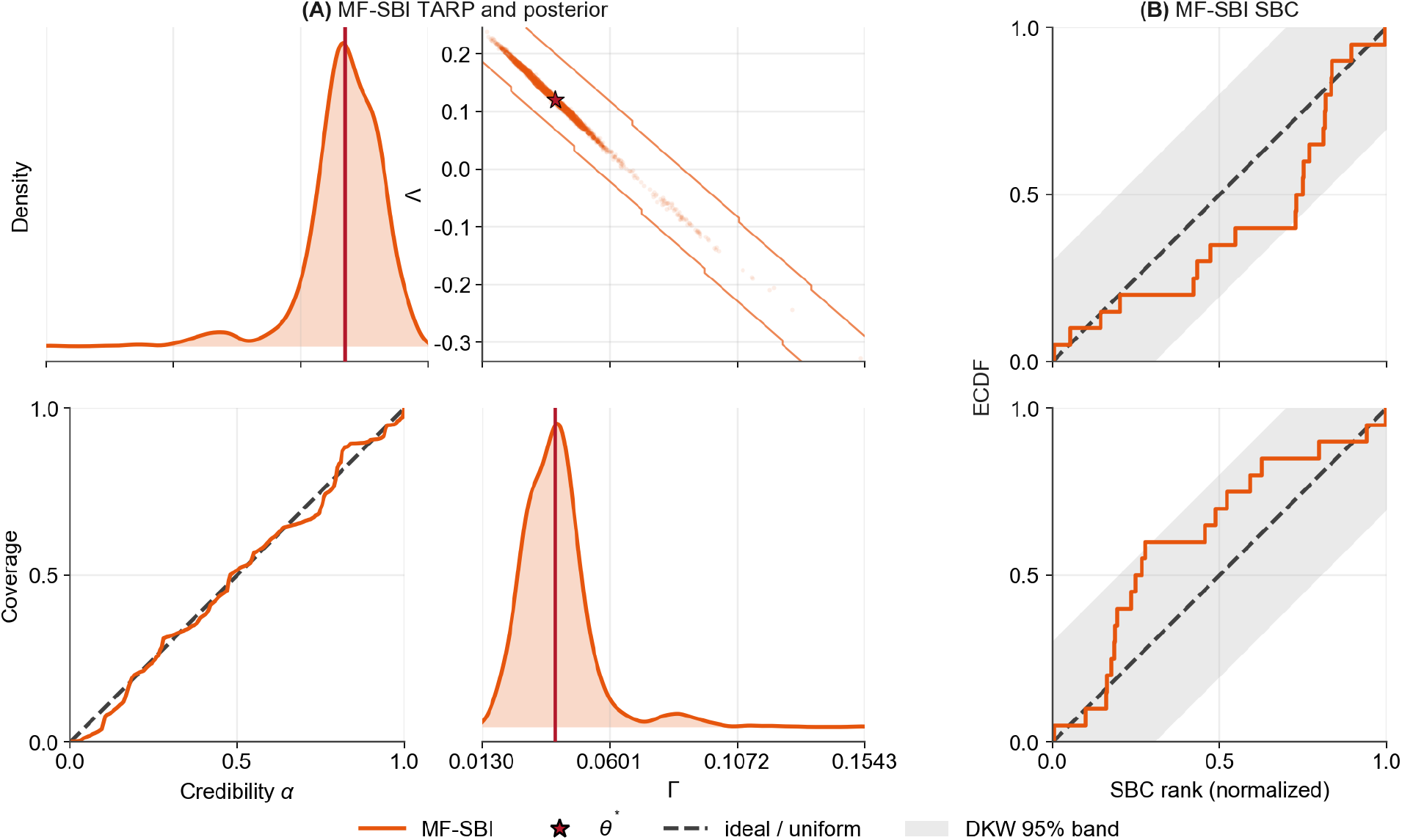
Calibration of MF-SBI with the. Δ*T*_0.05_ **LF simulator** (10000 LF and 500 HF simulations). (A) Marginal and joint posteriors for the primary observation, with TARP expected coverage against credibility level over 20 test observations; the dashed line shows perfect calibration. (B) Empirical cumulative distribution functions of normalised SBC ranks for Λ and Γ over the same observations, with 95% Dvoretzky–Kiefer–Wolfowitz bands (grey).

For the primary observation, the posterior was centred on *θ*^*^ (Fig 5A). Across the 20 test observations, TARP coverage stayed close to the line expected for a well-calibrated posterior (Fig 5A), and the distributions of SBC ranks for Λ and Γ tracked the uniform ECDF and largely remained within the 95confidence bands, with only small local excursions (Fig 5B), with mean normalised ranks of 0.61 and 0.37 respectively, close to the value of 0.5 expected under correct calibration. These results indicate that credible intervals from this configuration contain the true parameters at close to their stated rate. With 20 test observations, these checks can detect substantial but not subtle miscalibrations.

### The same pattern holds for a second vertex model fitted to imaging data

To test whether these findings depend on the particular model and synthetic observation used so far, we repeated screening and inference on a second vertex model, which has a different anisotropic mechanical description and no cell division, using an observation computed from 23 published images of the *Drosophila* pupal wing at 16.5–18.5 h after puparium formation [9] (*Materials and methods*). Posterior accuracy is evaluated at 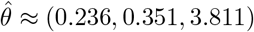 , the high-fidelity parameter whose summary statistics lie closest to the observation (Table 4). All comparisons were repeated five times.

The same three groups of candidate emerged (Fig 6A). Candidates using a larger timestep (Δ*T*_2_, Δ*T*_5_, Δ*T*_10_) or stopping at half the HF end time (*T*_50%_) were cheap and accurate; those stopping earlier (*T*_12%_, *T*_25%_), using a smaller tissue (*M*_4_) or a regular hexagonal initial condition (Honey) were inaccurate; and those averaging fewer realisations (*K*_1_, *K*_2_, *K*_5_) or using a smoothed initial condition (Voro+5) were accurate but costly. Notably, shortening the simulation was safe for this model provided it was not shortened too much, unlike for the proliferating model. This is consistent with the absence of cell division: the summary statistics depend only on how far the tissue has relaxed, rather than on a sequence of divisions that must be completed. Which approximations are safe therefore depends on the model, and cannot be assumed in advance.

**Fig 6.**
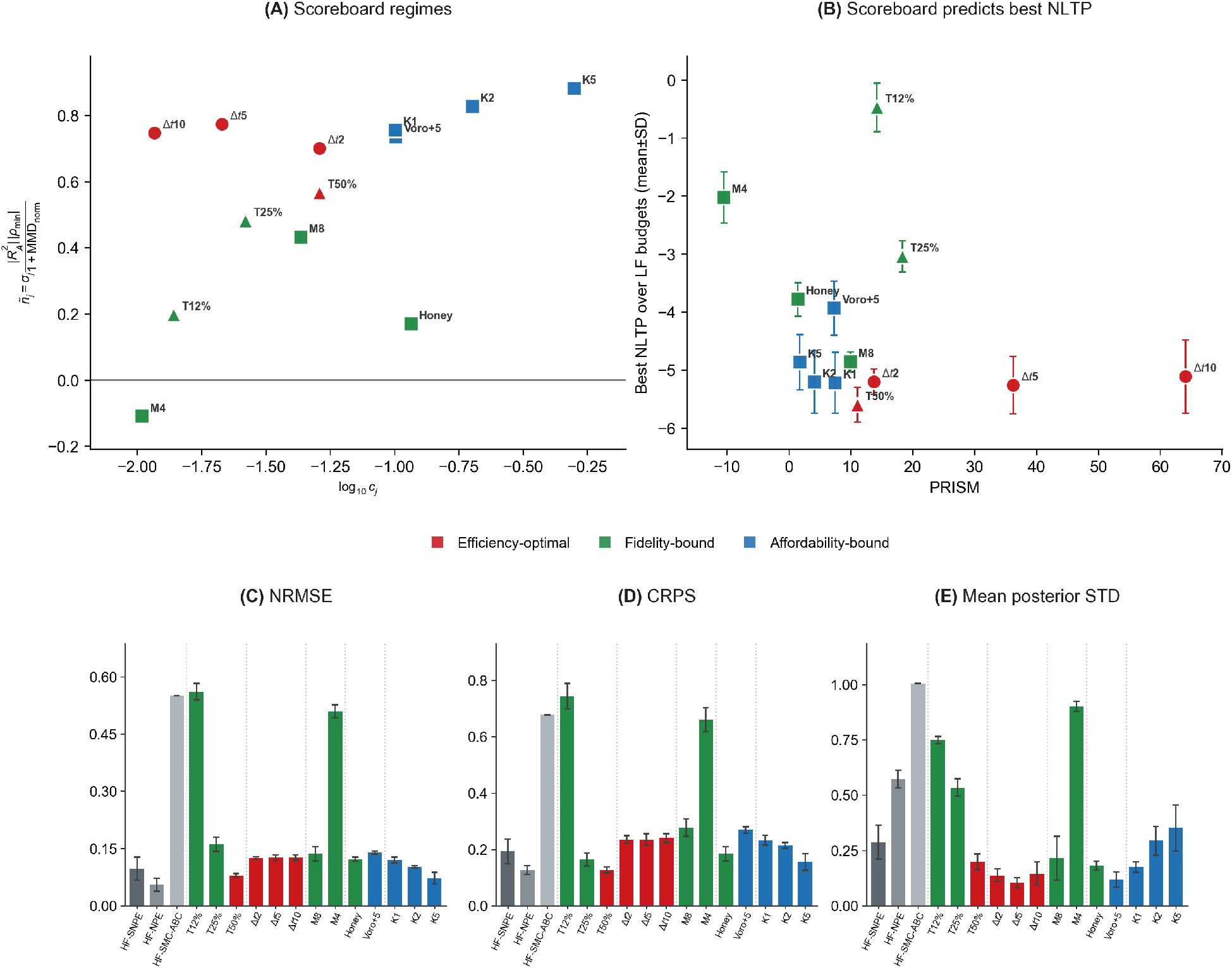
Screening and inference for the vertex model fitted to pupal-wing imaging data. (A) Accuracy term of the PRISM score plotted against relative cost for twelve LF candidates, separating efficiency-optimal (red), fidelity-bound (green), and affordability-bound (blue) candidates. (B) Best NLTP after MF-SBI training (mean *±* standard deviation (SD) over five repeats) against PRISM score. (C–E) NRMSE, CRPS, and mean posterior SD for each method at the largest affordable LF budget. All measures are evaluated at the data-matched reference 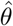 .

As for the proliferating model, candidates with higher PRISM scores tended to give lower NLTP after training (Fig 6B). The uniform prior has NLTP *≈* 0.40. Over five repeats, the median NLTP was *−* 0.45 for SMC-ABC, from a Gaussian kernel density on its 300 particles, 0.10 for NPE, and 26 for SNPE. The median is reported because one SNPE repeat lies far from the others. The efficiency-optimal candidates reached much lower values: the median of NLTP was about *−*5.7 for *T*_50%_ and about *−*4.6 to *−*4.8 for the three coarsened timesteps. On NRMSE, CRPS and posterior width, the efficiency-optimal candidates performed comparably to NPE and SNPE, while the fidelity-bound candidates and SMC-ABC performed worst (Fig 6C–E). SNPE gave one of the narrowest posteriors but a poor NLTP, showing that a narrow posterior can indicate overconfidence rather than accurate recovery of the reference parameters.

Pairwise posteriors for the three parameters show the same pattern (Fig 7). SMC-ABC gave broad posteriors with little probability near 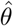 , while NPE and SNPE gave narrow posteriors that were not centred on 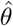. MF-SBI with *T*_50%_ gave compact posteriors close to 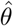 for every pair of parameters. Posteriors for all twelve candidates are shown in S2 Fig. For this observation and budget, MF-SBI with a screened LF simulator therefore recovered the data-matched reference more reliably than inference using HF simulations alone.

**Fig 7.**
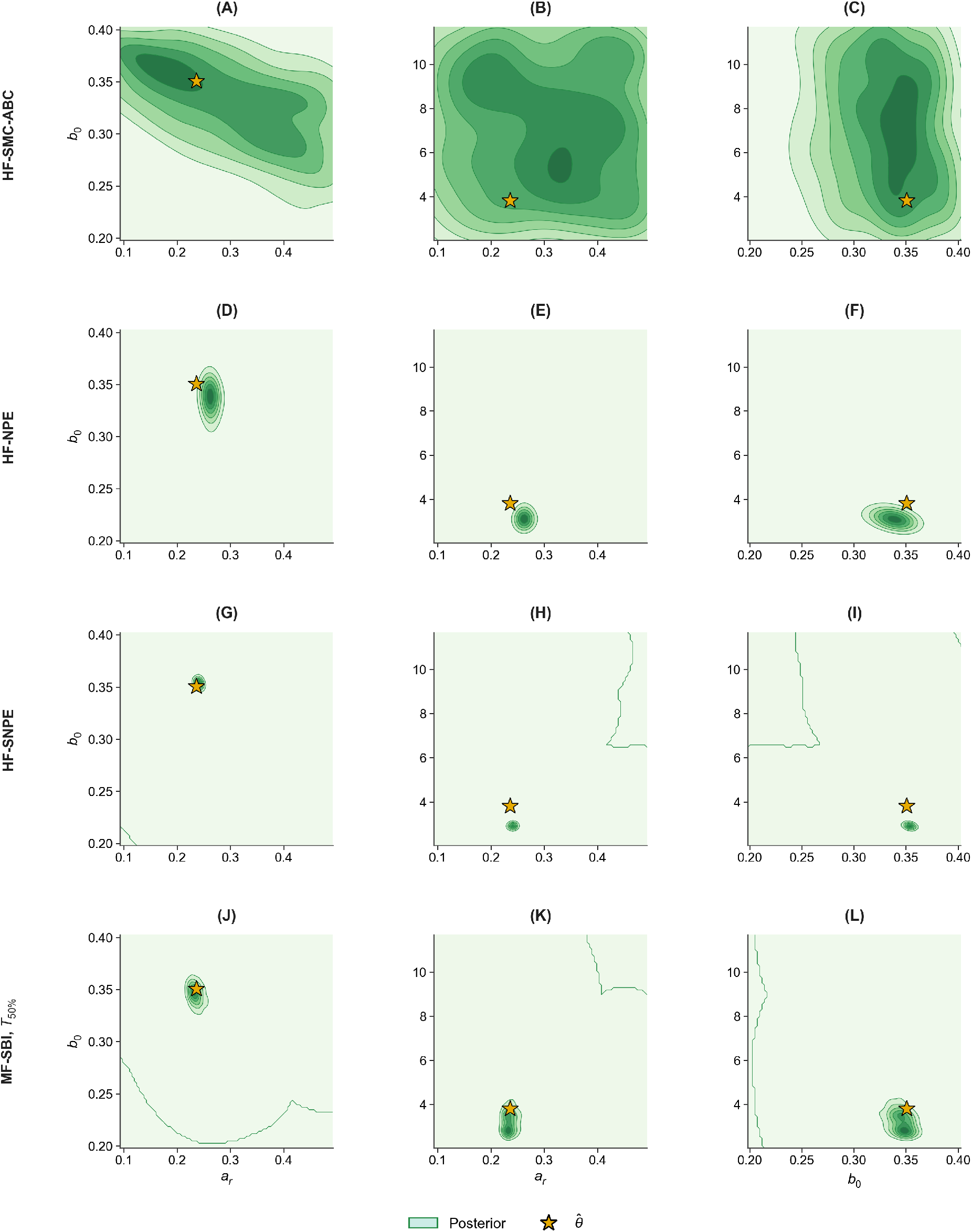
Posteriors for the pupal-wing observation at a matched budget. Pairwise posterior samples for (*a*_*r*_, *b*_0_, *k*) from (A) SMC-ABC, (B) NPE, and (C) SNPE, each using HF simulations only, and (D) MF-SBI with the *T*_50%_ LF simulator. Stars mark 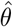 , the high-fidelity parameter matched to the observation.

## Discussion

Multi-fidelity simulation-based inference offers a way to calibrate computationally expensive stochastic models, such as vertex models of proliferating tissues, at a fraction of the usual cost. However, existing methods assume that a suitable low-fidelity simulator has already been chosen, whereas in practice several cheaper versions of a simulator can usually be constructed, and these are not interchangeable. In this work, we have introduced PRISM, a screening procedure that compares candidate LF simulators with the HF simulator using paired simulations, before any neural network is trained. By weighing agreement in summary statistics against relative cost, PRISM separates candidates that are cheap and accurate from those that are accurate but too costly, and from those that are inaccurate regardless of cost. Rather than selecting a single best candidate, PRISM identifies the group of candidates worth training, leaving measures of posterior accuracy to select among them. In weighing agreement against cost, PRISM follows the same principle as model selection in multifidelity Monte Carlo estimation [32] and the choice of continuation probabilities in multifidelity ABC [33], but applies it before training to the summary statistics used for inference, and adds a measure of distributional agreement, since a neural density estimator pretrained on LF simulations can be misled by systematic differences that correlation alone does not detect.

For the vertex model of a proliferating tissue, we found that the groups identified by screening matched the outcome of inference. Using a larger timestep preserved close agreement with the HF summary statistics at a fraction of the cost, and went on to give the most accurate posteriors of any method. Averaging over fewer realisations also preserved agreement, but each simulation still covered the full duration at the HF timestep; consequently, too few could be afforded to be useful. Shortening the simulation duration gave the poorest agreement and the least accurate posteriors, in one case placing less probability on the true parameters than the prior. This failure has a simple explanation: simulations stopped before proliferation is complete, and the tissue has relaxed, do not yet show the cell packing that the summary statistics describe. More generally, our results show that an LF simulator need not reproduce every aspect of the HF simulator, but must preserve the features of the output on which inference relies, at a cost low enough to allow many simulations. Correlation with the HF simulator alone was not sufficient: the intermediate duration *T*_350_ was positively correlated with the HF simulator in every summary statistic, yet still gave a poor posterior.

It is interesting to compare these findings with our earlier work on the same model. We previously showed that the choice of timestep affects quantitative predictions of vertex models [13], and that reliable parameter estimates require summary statistics averaged over repeated simulations [14]. The present results suggest that, for inference based on cell-packing statistics, moderate timestep coarsening introduces errors small enough to be corrected using a modest number of HF simulations, whereas the need for repeated simulations remains. Indeed, the HF outputs used in MF-SBI are themselves averages over ten realisations. Under a matched budget equivalent to 1,000 HF simulations, MF-SBI with a screened LF simulator gave more accurate posteriors than SMC-ABC, NPE and SNPE, and its posteriors were well calibrated across the test observations we considered.

For the second vertex model, fitted to imaging data from the *Drosophila* pupal wing, the same three groups of candidate emerged, and MF-SBI with a screened LF simulator again recovered the reference parameters more reliably than HF-only inference. However, the details differed. Halving the simulation duration was safe for this model, which has no cell division, whereas shortening the duration was the least safe approximation for the proliferating model. Which approximations are safe therefore depends on the model, and on the summary statistics used for inference, and cannot be assumed in advance. This is the main reason we believe that a screening step of this kind should precede multi-fidelity inference.

Our study has several limitations. First, accuracy comparisons for the proliferating model are based on a single synthetic observation, and calibration was assessed only for the selected configuration and with 20 test observations, which can reveal substantial but not subtle miscalibration. Second, the pupal-wing images are of a real epithelium. The method comparison is made at 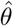, the high-fidelity parameter whose summaries match that observation. Those images were not generated by the vertex model. Third, the simulations used for screening are not counted against the inference budget. At the relative cost of each simulator, and including every screening run, the proliferating screen costs about 230 high-fidelity units and the pupal-wing screen about 260. In each screen, 20 of those units are the high-fidelity runs.. This cost is incurred once per model and set of candidates, and could be shared across many observations, but it should be taken into account when the budget is small. Fourth, each candidate changed either the duration or the timestep of a single realisation, or the number of realisations averaged, but we did not screen other combinations, such as a larger timestep averaged over several realisations, or a shorter duration combined with a larger timestep. Fifth, inference used the six summary statistics of cell packing from our previous work [14]; other statistics, or features learned directly from segmented images, may change both the screening results and the posteriors that follow. Finally, because neural density estimators are trained by stochastic optimisation, the posterior obtained with a given LF simulator cannot be predicted exactly without training. The screen can therefore identify which candidates are worth training, but not rank them precisely, as illustrated by the reversed order of Δ*T*_0.10_ and Δ*T*_0.05_.

Several extensions follow naturally from these limitations. Combined LF simulators could be screened in the same way, to test whether approximations that are individually safe remain so when combined. Simulation failures, which were common for several candidates (Fig 8), could be incorporated into the score rather than reported separately. The screening criteria could also be revisited for richer summary statistics. More broadly, the question of which cheaper approximations preserve the information needed for inference is not specific to vertex models. Cellular Potts, agent-based, and spatial-stochastic models of tissues admit their own reduced-cost versions, for example using coarser spatial grids, fewer agents or simplified reaction kinetics, and since PRISM requires only paired simulations and their summary statistics, it can be applied to these models without modification. To support this, our implementation is openly available (see Software and reproducibility).

**Fig 8.**
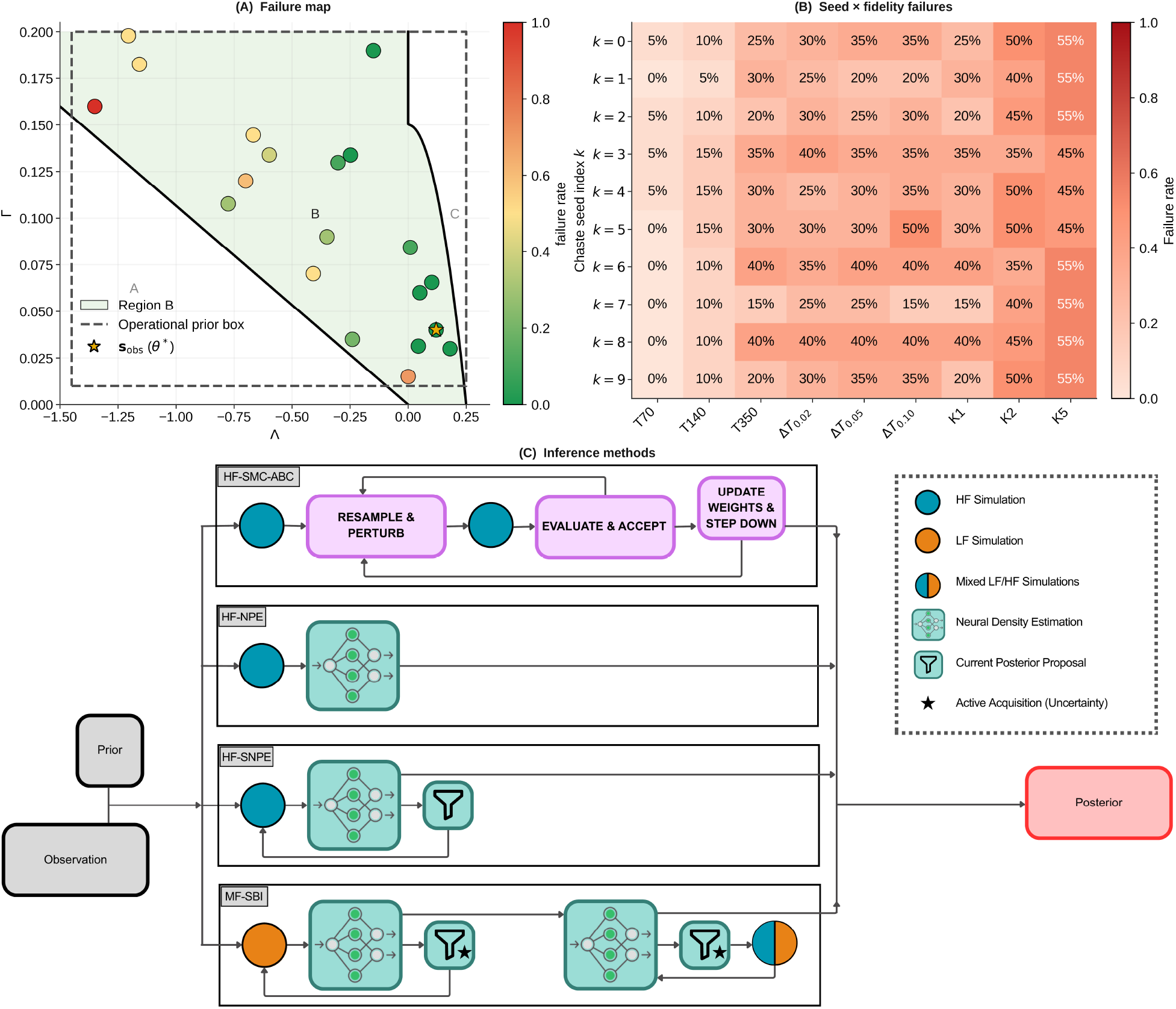
Operational prior, simulation failures, and inference allocation for the proliferating model. (A) Regions A–C in (Λ, Γ) with the synthetic observation *θ*^*^ and HF failure rates at the *N*_*θ*_ = 20 screening locations (*K* = 10 matched seeds). (B) Failure rates by matched seed across the nine LF candidates. (C) HF-SMC-ABC, HF-NPE, HF-SNPE, and MF-SBI paths from prior and observation to posterior, contrasting HF-only, LF, and mixed LF/HF allocation.

In conclusion, multi-fidelity inference of a stochastic tissue model is only as reliable as the LF simulator used for most of the computation. By screening candidate simulators before training, PRISM identifies those that are cheap enough to be useful and accurate enough to be trusted. In doing so, it makes Bayesian inference of mechanical parameters from tissue data feasible at a modest computational cost, opening the way to comparing tissue mechanics across genotypes, developmental stages or experimental conditions.

## Materials and methods

### Overview

Our workflow has three stages (Fig 1). First, we construct a set of candidate LF simulators and screen them with PRISM, using paired LF-HF simulations and no neural network training. Second, we perform MF inference with each candidate and compare the resulting posteriors with those from three single-fidelity (HF-only) baselines under a matched computational budget. Third, for the configuration selected on the basis of posterior accuracy, we check posterior calibration across a set of independent test observations.

Screening is carried out before, and independently of, inference, so that the PRISM score assigned to a candidate cannot be influenced by how well it later trains. We apply this workflow to two vertex models: a model of a proliferating tissue, which provides a synthetic test case with known ground truth; and a model of a non-proliferating tissue fitted to published imaging data. Table 1 summarises how the present study relates to our previous ABC analysis of the former model [14].

**Table 1.**
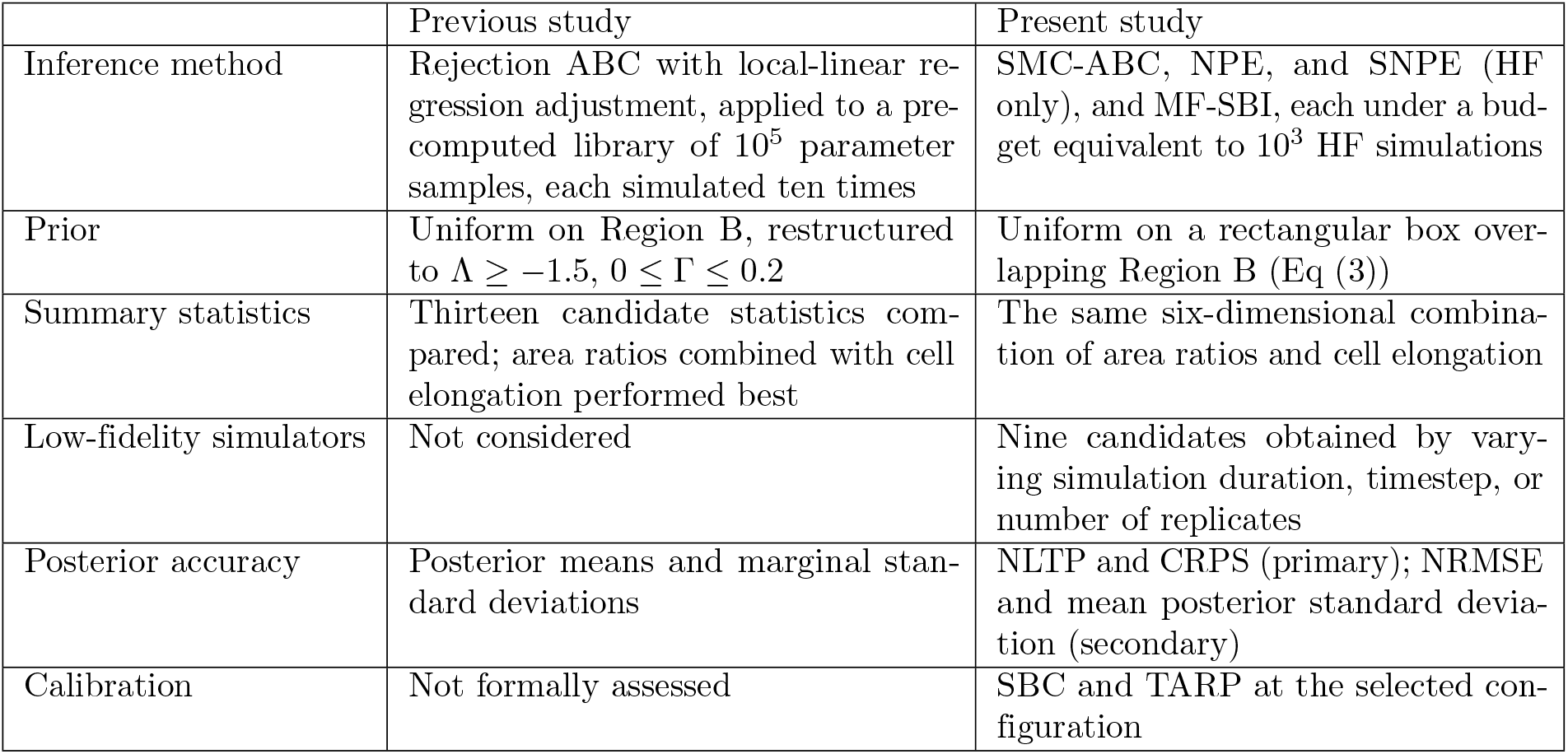
Comparison of our previous ABC analysis of the vertex model of a proliferating tissue with the present study.

|  | Previous study | Present study |
|---|---|---|
| Inference method | Rejection ABC with local-linear re-gression adjustment, applied to a pre-computed library of 10<sup>5</sup> parameter samples, each simulated ten times | SMC-ABC, NPE, and SNPE (HF only), and MF-SBI, each under a bud-get equivalent to 10<sup>3</sup> HF simulations |
| Prior | Uniform on Region B, restructured to Λ ≥ −1.5, 0 ≤ Γ ≤ 0.2 | Uniform on a rectangular box over-lapping Region B (Eq (3)) |
| Summary statistics | Thirteen candidate statistics com-pared; area ratios combined with cell elongation performed best | The same six-dimensional combina-tion of area ratios and cell elongation |
| Low-fidelity simulators | Not considered | Nine candidates obtained by vary-ing simulation duration, timestep, or number of replicates |
| Posterior accuracy | Posterior means and marginal stan-dard deviations | NLTP and CRPS (primary); NRMSE and mean posterior standard devia-tion (secondary) |
| Calibration | Not formally assessed | SBC and TARP at the selected con-figuration |

### Vertex model of a proliferating tissue

We use the vertex model of a growing *Drosophila* wing imaginal disc analysed in our previous work [14], implemented in Chaste [11]. The apical surface of the epithelium is represented as a tessellation of polygons, each corresponding to a cell, whose vertices move according to overdamped dynamics,

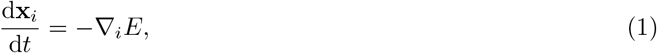

where **x**_*i*_ is the position of vertex *i* and _*i*_ denotes the gradient with respect to **x**_*i*_. Following Farhadifar and colleagues [12], the non-dimensionalised energy of the tissue is

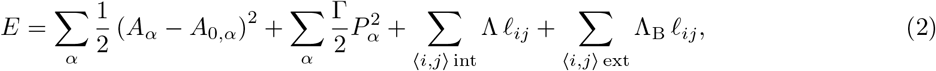

where *A*_*α*_, *A*_0,*α*_ and *P*_*α*_ denote the area, target area, and perimeter of cell *α*, and *l*_*ij*_ denotes the length of the edge joining vertices *i* and *j*. The first two sums run over all cells, the third over edges in the interior of the tissue, and the fourth over edges on the tissue boundary. Space has been rescaled by a characteristic cell length and time by the mechanical relaxation timescale, so that the area elasticity coefficient is one [14]. The two parameters we infer are the line tension, Λ, which captures the combined effect of adhesion and contractility along cell-cell junctions, and Γ, which captures the contractility of each cell’s actomyosin cortex. To prevent unphysical tissue shapes at the free boundary, we set the boundary line tension to Λ_B_ = Λ if Λ *≥* 0 and Λ_B_ = 0 otherwise [14].

Each simulation begins with four hexagonal cells of unit area and proceeds through *n*_d_ = 7 rounds of cell division. Each cell cycle comprises an interphase, whose duration is drawn independently from an exponential distribution with mean 2*t*_cycle_*/*3, followed by a growth phase of fixed duration *t*_cycle_*/*3, during which the cell’s target area increases linearly from one to two. The cell then divides along its short axis, and each daughter cell is assigned unit target area. Cells of the final generation remain in interphase for the rest of the simulation. The random interphase durations are the only source of stochasticity in the model. With *t*_cycle_ = 20, the seven rounds of division take on average around 140 time units, after which the tissue, comprising approximately 500 cells, relaxes to a static configuration by the end time *T*_end_ = 700. Cell neighbour exchanges (T1 transitions) occur whenever an edge becomes shorter than *l*_T1_ = 0.01, and small triangular cells are removed (T2 transitions) whenever their area falls below *A*_T2_ = 0.001. In oour high-fidelity runs at *θ*^*^, both transitions occurred. Overlapping boundary cells are merged as described previously [14]. Eq (1) is solved using a forward Euler scheme with timestep Δ*T*. All model parameter values are listed in Table 2.

**Table 2.** Parameter values for the vertex model of a proliferating tissue (non-dimensionalised). Values follow [14] unless stated otherwise.

| Parameter | Description | Value |
|---|---|---|
| Λ | Line tension | Inferred |
| Γ | Perimeter contractility | Inferred |
| Λ<sub>B</sub> | Boundary line tension | max(Λ, 0) |
| *A*<sub>0</sub> | Target area in interphase | 1 |
| *t*<sub>cycle</sub> | Mean cell-cycle duration | 20 |
| *n*<sub>d</sub> | Number of rounds of division | 7 |
| *l*<sub>T1</sub> | T1 transition threshold | 0.01 |
| *ρ* | Ratio of new to threshold edge length after a T1 transition | 1.5 |
| *A*<sub>T2</sub> | T2 transition area threshold | 0.001 |
| Δ*T* | Timestep (HF) | 0.01 |
| *T*<sub>end</sub> | Simulation end time (HF) | 700 |

#### Summary statistics

As in our previous work [14], we summarise each simulated tissue by six statistics of cell packing, which gave the most accurate parameter estimates among those we tested: the mean area of cells with *n* sides, divided by the mean area of all cells, for *n* = 4, … , 8 (‘area ratios’), and the mean cell elongation. The elongation of a cell is defined as the square root of the ratio of the largest to the smallest eigenvalue of the moment of inertia tensor of the polygon representing it. All statistics are computed over cells that do not share an edge with the tissue boundary, since cells at the boundary have different shapes from those in the bulk of the tissue.

#### Parameter space and prior

Not every combination of (Λ, Γ) gives rise to a physically meaningful tissue. Previous analyses of the energy in Eq (2) [12, 41] divide parameter space into three regions (Fig 8A). In Region A, the energy has no unique minimum and simulated tissues tend to develop self-intersecting cells; in Region B, the energy has well-defined minima and cell packing varies with the parameters; and in Region C, cells shrink towards zero area and are progressively removed. In our previous work we restricted the prior to Region B [14]. Here the prior is a rectangle. It contains Region B as drawn in Fig 8A and also parts of Regions A and C, and every simulator draws its parameters from it. High-fidelity runs fail inside Region B, more often at smaller Λ (Fig 8A), and the low-fidelity candidates fail at different rates (Fig 8B). Restricting the prior to Region B would treat those simulators as if they shared one high-fidelity failure map, whereas using a larger box allows us to capture and compare the low-fidelity choices and their failures inside and outside Region B (SI Fig. 1). we instead use the uniform prior

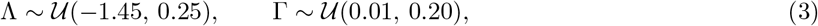

Some simulations fail to complete, for example through the formation of self-intersecting cells, numerical instability, or exceeding a wall-clock limit. A simulation is stopped after 3600 seconds, or after 900 seconds without progress. Such failures depend jointly on the parameters, the simulator settings, and the random seed (Fig 8A,B), and we treat them as part of the behaviour of the forward model. For NPE, SNPE, and MF-SBI, a failed simulation is left out of the training set and still counts toward the budget. SMC-ABC counts the call in the same way and returns a large constant summary, which the distance then rejects.

#### High-fidelity simulator and observations

Our HF simulator uses *T*_end_ = 700 and Δ*T* = 0.01, as in [14]. Since summary statistics computed from a single tissue give poorly identifiable posteriors [14], each HF output is the mean of the summary statistics of *K* = 10 independent realisations of the model. Throughout, one ‘HF simulation’ refers to one such ten-realisation output, and computational costs are expressed in these HF-equivalent units. The primary synthetic observation, **s**_obs_, is a single HF output at *θ*^*^ = (0.12, 0.04), a parameter set commonly used in vertex model simulations of the wing disc [12]. For the calibration analysis we used 20 of the parameters at which all ten high-fidelity replicates finished (Fig 5). We drew 200 parameters uniformly from the prior box and ran ten high-fidelity replicates at each. All ten replicates finished for 40 of those parameters, and the 20 calibration observations are 20 of those 40.

### Candidate low-fidelity simulators

We constructed nine LF simulators from the HF simulator (Table 3). Six use a single realisation (*K* = 1) with either a shorter simulation duration (*T*_end_ ∈ {70, 140, 350}) or a larger timestep (Δ*T* ∈ {0.02, 0.05, 0.10}), with all other settings as in the HF simulator. Given the time taken for seven rounds of division (see above), the shortest duration stops simulations part-way through proliferation. The remaining three use HF settings but average over fewer realisations (*K* ∈ {1, 2, 5}). Note that *K*_1_ is therefore the single-realisation counterpart of the HF simulator against which the duration and timestep candidates differ in one further setting. We define the relative cost *c*_*j*_ of candidate *j* as the ratio of its median wall-clock time to that of the HF simulator, measured on the same hardware. The median is taken over completed runs among those 200 runs we used for LF screening, and failed runs are not considered.

**Table 3.**
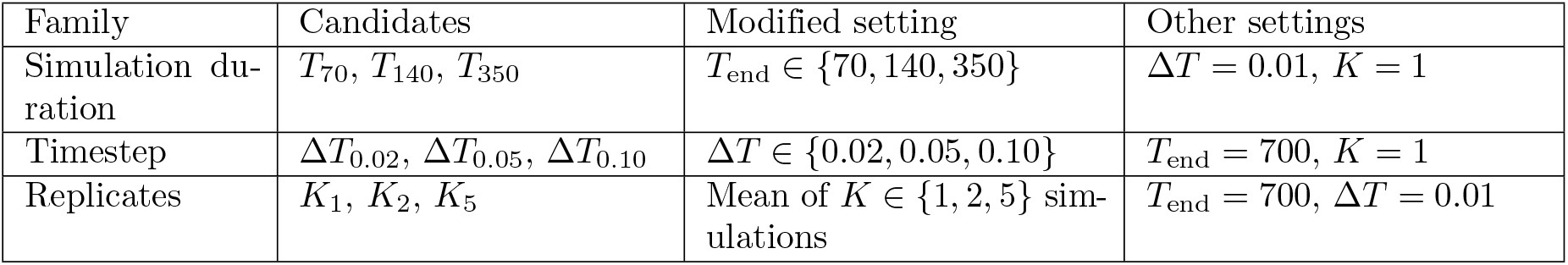
Candidate LF simulators for the model of proliferating epithelial tissue. The HF simulator has *T*_end_ = 700, Δ*T* = 0.01, and *K* = 10.

| Family | Candidates | Modified setting | Other settings |
|---|---|---|---|
| Simulation du-ration | T<sub>70</sub>, T<sub>140</sub>, T<sub>350</sub> | T<sub>end</sub> ∈ {70, 140, 350} | ΔT = 0.01, K = 1 |
| Timestep | ΔT<sub>0.02</sub>, ΔT<sub>0.05</sub>, ΔT<sub>0.10</sub> | ΔT ∈ {0.02, 0.05, 0.10} | T<sub>end</sub> = 700, K = 1 |
| Replicates | K<sub>1</sub>, K<sub>2</sub>, K<sub>5</sub> | Mean of K ∈ {1, 2, 5} sim-ulations | T<sub>end</sub> = 700, ΔT = 0.01 |

### Vertex model fitted to *Drosophila* pupal-wing imaging data

To test the approach on a different model and on experimental data, we implemented a second vertex model in Chaste, based on the anisotropic epithelial mechanics used by Ogita, Yan and colleagues to analyse the *Drosophila* pupal wing [8, 9]. Those studies inferred parameters by fitting a force-balance condition to fixed, segmented cell outlines. Since PRISM and MF-SBI require a forward simulator, we instead simulate a short mechanical relaxation of the tissue, without cell division. The tension on an internal edge of length *l* and orientation *φ* relative to the proximal-distal axis is

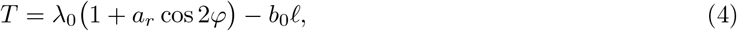

with *λ*_0_ = 1, and the pressure in a cell of area *A* is *P* = *−k*(*A − A*_0_), with *A*_0_ = 1. Vertices move according to overdamped dynamics driven by these tensions and pressures. We infer *θ* = (*a*_*r*_, *b*_0_, *k*), which respectively set the anisotropy of junctional tension, its dependence on edge length and the stiffness of cell area, with independent uniform priors

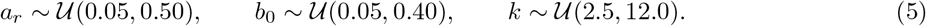

Each simulation begins from a random Voronoi tessellation with *N* = 144 cells, chosen to match the number of internal cells in the imaging data (typically 84–179 per image), and relaxes until *T*_end_ = 5 with timestep Δ*T* = 0.001. The 144 points are drawn independently from the uniform distribution on a square, and the mesh is their Voronoi tessellation, scaled to unit mean cell area, with no iteration of Lloyd’s algorithm Lloyd’s algorithm [42]. Neighbour exchanges occur when an edge is shorter than 0.10, and a triangular cell is removed when its area falls below 0.001. In twenty high-fidelity runs, each at a different draw from the prior, neighbour exchanges occurred in every run, 74 to 2,974 of them during the relaxation, and a triangular cell was removed in two of the twenty. The random initial tessellation is the source of stochasticity. Each high-fidelity output is the mean of *K* = 10 such realisations.

### Observation

The observation is computed from 23 segmented images of the pupal wing at 16.5–18.5 h after puparium formation, published by Yan and colleagues [9]. The images and their segmentation are those described by Ogita and colleagues [8]: adherens-junction outlines of the wing blade, with the hinge marked separately in their schematic, aligned to the proximal–distal axis. Vertices and their connectivity were taken from the skeletonized images. For each image, and for each simulated tissue, we compute five summary statistics: the mean and standard deviation of log edge length, the mean of cos 2*φ* over internal edges, the mean log cell area, and the fraction of hexagonal cells. These summaries are the geometric features controlled by the inferred parameters. *a*_*r*_ multiplies cos 2*φ* in the junction tension; therefore, the mean of cos 2*φ* measures how strongly the internal edges align with the proximal–distal axis. *φ* is the angle of an edge. Lengths and areas are scaled by the median area of the internal cells, in the same way for every image and every simulation. The scaled edge lengths are logged because the shortest edges are only about 5% of the median edge length, and the logarithm gives those short edges weight in the mean and the standard deviation. The observation **x**_*o*_ is the mean of these statistics over the 23 images (Table 4). Posterior accuracy is evaluated at 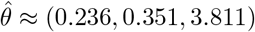, taken from the 3,000 high-fidelity simulations of SMC-ABC, NPE, and SNPE, 1,000 from each method. The HF summary statistics at 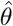 differ from the observation by about 2% of its norm, and by less than the image-to-image standard deviation in every component (Table 4).

**Table 4.**
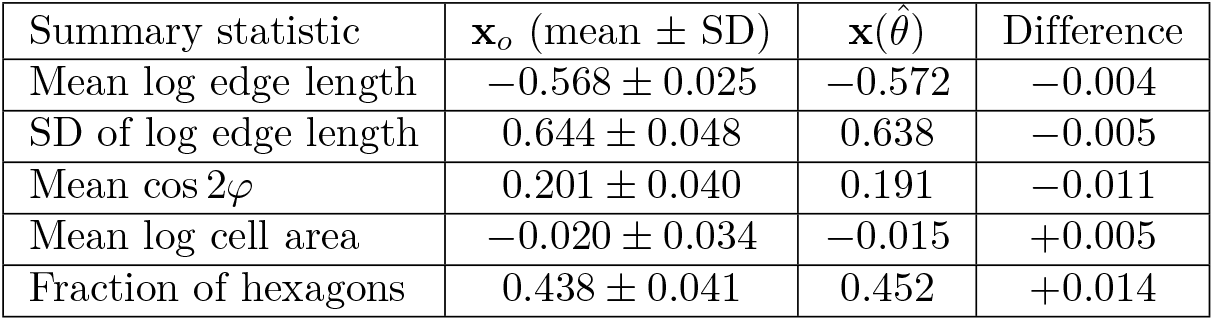
Agreement between the pupal-wing observation and HF summary statistics at the data-matched reference. The observation **x**_*o*_ is given as mean *±* standard deviation over 23 images; 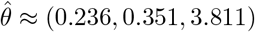 .

| Summary statistic | **x**<sub>*o*</sub> (mean ± SD) | **x**(θ̂) | Difference |
|---|---|---|---|
| Mean log edge length | −0.568 ± 0.025 | −0.572 | −0.004 |
| SD of log edge length | 0.644 ± 0.048 | 0.638 | −0.005 |
| Mean cos 2φ | 0.201 ± 0.040 | 0.191 | −0.011 |
| Mean log cell area | −0.020 ± 0.034 | −0.015 | +0.005 |
| Fraction of hexagons | 0.438 ± 0.041 | 0.452 | +0.014 |

### Candidate low-fidelity simulators

We constructed twelve LF candidates for this model. Three stop the relaxation early, at 12.5%, 25%, and 50% of the HF end time (*T*_12%_, *T*_25%_, *T*_50%_). Three use a timestep two, five, and ten times larger than the HF timestep (Δ*T*_2_, Δ*T*_5_, Δ*T*_10_). One uses a smaller tissue of 4 *×* 4 cells (*M*_4_). Two replace the random Voronoi initial condition, either with a regular hexagonal packing (Honey) or with a Voronoi tessellation smoothed by five iterations of Lloyd’s algorithm [42] (Voro+5). The remaining three use HF (*K*_10_) settings but average over one, two, or five realisations (*K*_1_, *K*_2_, *K*_5_). Screening, inference, and evaluation then follow the procedures described below for the model of a proliferating epithelial tissue, with only the model, parameters, summary statistics, and LF candidates changed. For this model the screen uses 20 parameter values in the prior box and ten seeds at each, and the inference budget is 1,000 high-fidelity units.

### PRISM screening

#### Paired simulations

PRISM compares each candidate LF simulator with the HF simulator using paired simulations, before any neural network is trained. For the model of a proliferating epithelial tissue, we chose *N*_*θ*_ = 20 parameter values in Region B, and at each we ran *K* = 10 realisations. These are independent uniform draws from the prior rectangle, kept only when they fell inside Region B. For each candidate *j* and each parameter-seed pair (*θ*_*i*_, *k*), we compared its summary statistics, 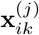 , with the HF output at the same parameter value, **y**_*i*_ (the mean over the ten realisations). Each such pair has one parent seed, shared by the high-fidelity run and every candidate. The cell-cycle seed of a realisation is derived from that parent seed, the candidate, and the realisation index. *K*_2_ averages two realisations drawn in this way, and *K*_5_ averages five, so those realisations are matched to the high-fidelity run at the same seed index and each has its own cell-cycle seed. Comparing with the ten-realisation HF mean matches the form of the HF outputs used for inference. The correlation, *R*^2^, and the discrepancy below are computed on these summaries on their original scales. Each summary component is standardised only inside the density estimator. Pairs for which either simulation failed are excluded, and failure rates are recorded separately (Fig 8B).

#### Screening metrics

No single statistic captures all the ways in which an LF simulator can differ from the HF simulator: LF and HF outputs may be highly correlated but systematically offset, close on average but differently distributed, or accurate but too costly. We therefore compute three measures of agreement over all valid pairs. The first is the smallest, over the six summary statistics, of the Pearson correlation coefficient between LF and HF values, *ρ*_min_(*j*) = min_*d*_ *ρ*_*d*_(*j*). We use the minimum rather than the mean so that strong agreement in some statistics cannot mask poor agreement in others. The second is the pooled coefficient of determination,

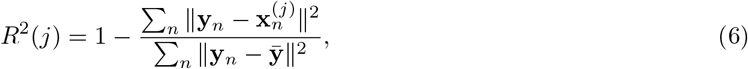

where *n* indexes valid pairs and 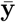 is the mean HF output. Unlike the correlation, *R*^2^ penalises systematic offsets, and it is negative when the LF outputs are further from the HF outputs than the HF mean is. The third is the normalised maximum mean discrepancy between the LF and HF output distributions, MMD_norm_(*j*), computed with a Gaussian RBF kernel whose bandwidth is the median of the positive pairwise squared distances of the pooled LF and HF outputs. That discrepancy is then divided by the mean of the squared norms of the HF outputs. It detects differences in distribution that the first two measures can miss. These screening metrics are computed on the raw summaries. Inside the density estimator, each summary component and each parameter is standardised separately.

#### RISM score

We combine these measures into an accuracy term and divide by the relative cost,

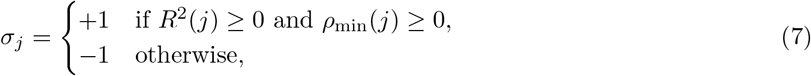

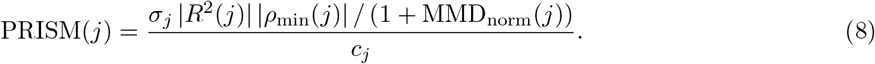

For candidates with *σ*_*j*_ = +1, the accuracy term lies between zero and one, taking the value one only if the candidate reproduces the HF outputs exactly. The sign *σ*_*j*_ ensures that a candidate failing either test receives a negative score, however cheap it is. Dividing by *c*_*j*_ gives PRISM units of accuracy per HF-equivalent simulation.

#### Reading the scoreboard

Rather than selecting a single winner by score, we plot the accuracy term against relative cost (Fig 3A), on which the PRISM score is the slope of the line from the origin to each candidate. Candidates then fall into three groups: cheap and accurate (‘efficiency-optimal’), accurate but costly (‘affordability-bound’), and inaccurate regardless of cost (‘fidelity-bound’). The groups are read from that plot. Relative cost differs across simulators and is the denominator of the PRISM score, so the plot shows which candidates are worth training. We carry the efficiency-optimal group forward to inference, and treat the ordering within it as indicative only. To relate cost to the inference budget, we also report the largest number of LF simulations that fits within a total budget of *B*_tot_ HF-equivalent units after *N*_HF_ HF simulations have been reserved,

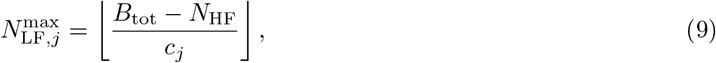

with *B*_tot_ = 1000 and *N*_HF_ = 500. This quantity is not used to assign groups. The procedure is summarised in Algorithm 1.

##### Algorithm 1

PRISM screening of candidate LF simulators.

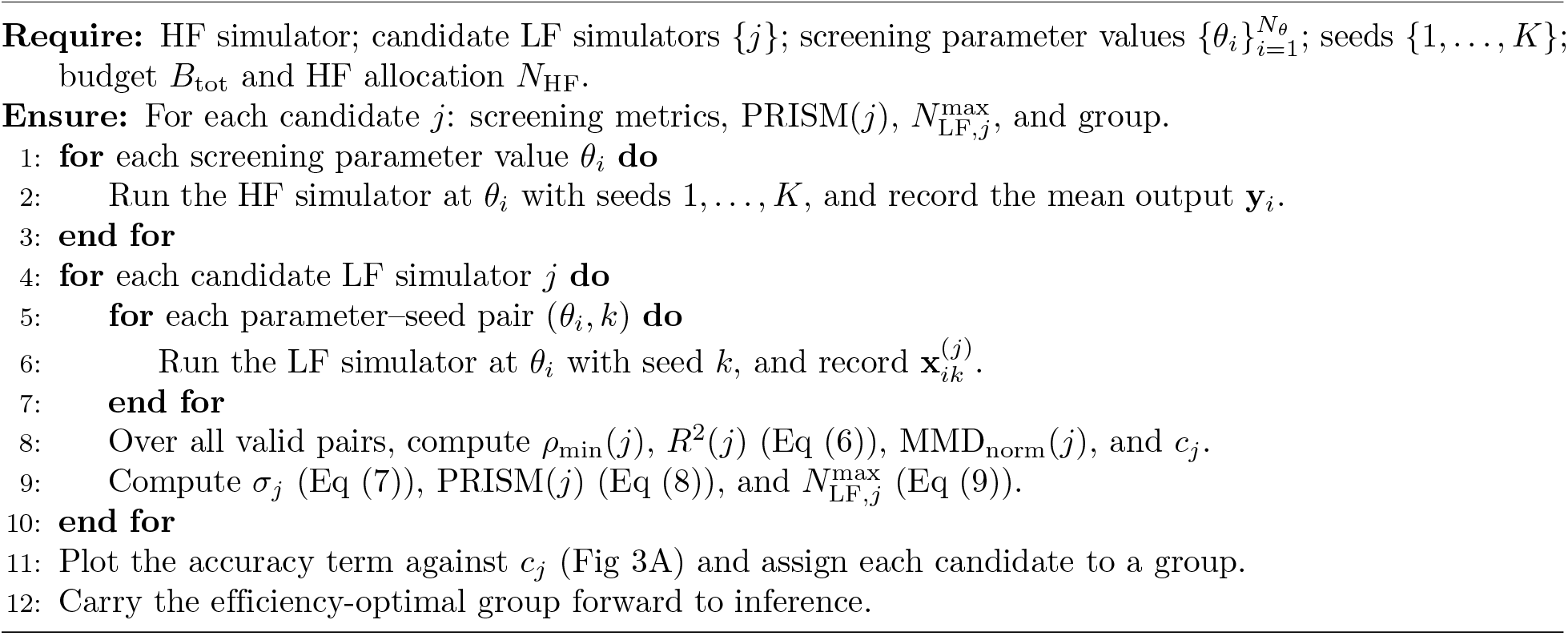

### Inference

#### Budget

All methods are given the same total budget of *B*_tot_ = 1000 HF-equivalent simulations. The HF-only baselines use 1000 HF simulations. For MF-SBI with candidate *j*, the budget is divided as *B*_tot_ = *N*_HF_ + *N*_LF_ *c*_*j*_ with *N*_HF_ = 500 and *N*_LF_ ∈{500, 1000, 5000, 10000}, capped at 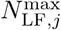. Training parameters are drawn from the prior (Eq (3) or Eq (5)), or from the current proposal for sequential methods. Each method is conditioned on the primary observation (for the proliferating model) or on **x**_*o*_ (for the pupal-wing model), and each configuration is repeated five times with different random seeds. A different seed produces different simulations. For an inference run it also changes the parameter draws and the initialisation of the density estimator. The same holds for the ten screening seeds and for the realisations inside a *K*_2_, *K*_5_, or *K* = 10 output.

### HF-SMC-ABC

Sequential Monte Carlo ABC [21, 22] maintains a population of weighted parameter values (particles), which it repeatedly resamples and perturbs while decreasing the tolerance on the distance between simulation and observed summary statistics. We use algorithm variant C with 300 particles, an L2 distance on the unscaled summaries, and a Gaussian perturbation kernel, and we stop when the simulation budget is reached. The next tolerance is the 0.2 quantile of the distances accepted in the previous population, and the Gaussian kernel uses half the weighted covariance of that population.

### HF-NPE

Neural posterior estimation [23] trains a conditional density estimator, *q*_*ϕ*_(*θ*| **s**), on pairs of parameter values drawn from the prior and the corresponding HF outputs, in a single round. The estimator is a neural spline flow with five transforms and 50 hidden units. Training uses Adam with learning rate 5 *×* 10^−4^, batches of 200, and a validation fraction of 0.1, and stops after 20 epochs without improvement in the validation loss, or at 200 epochs. We draw 2,000 samples from the trained posterior. Parameters are sampled from the prior and all HF simulations are pooled before optimisation, so the proposal is not adapted during training.

### HF-SNPE

Sequential neural posterior estimation uses the same density estimator and the same training schedule. We use SNPE-C, with five rounds of 200 simulations. The proposals stay on the prior support. After the first round, a fraction 0.9 of each new batch is drawn from the current posterior and a fraction 0.1 from the prior, and the training loss is the maximum-likelihood loss [26].

### MF-SBI

Our multi-fidelity procedure builds on the approach of Krouglova and colleagues [36], in which a density estimator is first trained on LF simulations and then refined using HF simulations. We allocate the simulations in three stages. Stage (i) uses half of the LF budget, drawn from the prior, to pretrain the density estimator, which is then copied to an ensemble of five. Stage (ii) uses one fifth of the LF budget, in two rounds. In each round, one fifth of the parameters are chosen from a pool of 1,000 prior draws as those with the largest variance, at the observation, of log *q*(*θ* | **s**) across the five estimators, and the rest of the round is drawn from the prior. Stage (iii) uses the remaining three tenths of the LF budget together with the 500 HF simulations, split over five rounds. Each round is drawn from the same posterior–prior mixture as SNPE, and the proposals stay on the prior support. Within a round the parameters are ranked by that variance: the highest-ranked share, in proportion to the HF simulations still unused, is run at high fidelity and the rest at low fidelity. The two outputs are pooled, and every network weight is updated on that pool. The full schedule is given in S1 Algorithm.

### Evaluating posteriors

No single measure fully characterises the quality of a posterior distribution, so we report several complementary measures. Unless stated otherwise, each is computed from 2000 posterior samples, and evaluated at *θ*^*^ for the model of proliferating epithelial tissue and at 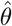 for the pupal-wing model.

#### Negative log target probability (NLTP)

Our primary measure is NLTP = log *q*(*θ*^*^ **s**_obs_), the negative log posterior density at the target. Lower values indicate that more posterior probability lies at the target, which is a stricter requirement than placing the posterior mean or mode near to it. An NLTP greater than that of the prior indicates a posterior that is less consistent with the target than the prior is. Since NLTP depends on the units of the parameters, we compare its values only within each model.

#### Continuous ranked probability score (CRPS)

For each parameter, we compute the CRPS [43], 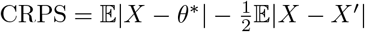 , where *X* and *X*^′^ are independent posterior samples, and average over parameters. That average is in the original parameter units, so the coordinate with the wider range contributes more. CRPS rewards posteriors that are both concentrated and centred on the target.

#### Normalised root mean square error (NRMSE)

We divide the error in each coordinate of the posterior mean by the width of that parameter’s prior, then take the root mean square. We report NRMSE as a secondary measure, since a posterior can have a mean close to the target while being broad, skewed, or displaced.

#### Posterior standard deviation

We report the mean of the marginal posterior standard deviation as a measure of posterior width. Like the CRPS average, it is left in the original parameter units. We interpret it only alongside NLTP and CRPS, since a narrow posterior is desirable only when it is centred on the target.

#### Calibration

For the configuration selected on the basis of the above measures (MF-SBI with Δ*T*_0.05_, *N*_LF_ = 10000, and *N*_HF_ = 500), we assess whether the posterior’s stated uncertainty is reliable, using simulation-based calibration (SBC) [39] and tests of accuracy with random points (TARP) [40]. For SBC, we compute the normalised rank of each true parameter value among 2000 posterior samples, for each of the 20 test observations, and compare the empirical cumulative distribution of these ranks with the uniform distribution expected under correct calibration, using 95% Dvoretzky-Kiefer-Wolfowitz confidence bands. TARP uses the same 20 test observations as SBC. Following Lemos et al., for each observation we draw 400 reference points uniformly in the rectangle spanned by those 20 parameter values and compare expected and observed coverage of the resulting credible regions.

### Software and reproducibility

Tissue simulations used Chaste from the develop branch at commit 0cff083, compiled with GCC 13.3.0. Inference was implemented in Python 3.12.3 with PyTorch 2.5.1 and sbi 0.25.0. Random seeds were fixed hierarchically for the paired screening simulations and for the inference experiments. Large simulation campaigns were run on CPU nodes of the University of Sheffield Stanage cluster. The general CPU nodes are Dell PowerEdge R650 machines with two 32-core Intel Xeon Platinum 8358 processors. The jobs requested CPUs and memory only. All code is available at https://github.com/maminium/prism-mf-sbi. The simulation outputs and the processed summaries are archived on Zenodo.

## Acknowledgments

We thank Min Tao and Bin Chen for helpful discussions on multi-fidelity approaches for vertex models during the BBSRC-funded project of which this work forms part. This study made use of the University of Sheffield’s Stanage high-performance computing cluster.

